# Alterations of the rewarding actions of amphetamine by prior nicotine and alcohol treatment: The role of age and dopamine

**DOI:** 10.1101/2020.08.22.263012

**Authors:** Andrea Stojakovic, Syed Muzzammil Ahmad, Kabirullah Lutfy

## Abstract

**Rationale:** Nicotine and alcohol each can serve as the gateway to other drugs.

**Objective:** The current study was sought to determine if prior nicotine and alcohol exposure alters amphetamine reward and if age and dopaminergic neurotransmission are involved.

**Methods:** Male and female adolescent and adult C57BL/6J mice were tested for baseline place preference, received six conditioning with saline/nicotine (0.25 mg/kg) twice daily followed by six conditioning with saline/ethanol (2 g/kg) in a counterbalance manner. Control mice were conditioned with saline/saline throughout. Finally, mice were conditioned with amphetamine (3 mg/kg) once in the nicotine-alcohol-paired chamber and then tested for CPP 24 h later. The following day, mice were challenged with amphetamine (1 mg/kg) and tested for CPP under a drugged state. Mice were then immediately euthanized, brain removed and nucleus accumbens isolated and processed for the expression of dopamine receptors and transporter, and glutamate receptors.

**Results:** We observed a greater amphetamine-induced CPP in adolescent than adult mice but no change in state-dependent CPP between the two age groups. In contrast, amphetamine-induced CPP in mice with prior nicotine-alcohol exposure was greater in adult than adolescent mice under both drug-free and drugged states. The enhanced response in adult mice was associated with greater expression of dopamine-transporter, reduced D2 receptors, and increased D1 receptors with no changes in glutamate receptors.

**Conclusions:** These results suggest that prior nicotine and alcohol exposure differentially alters the rewarding action of amphetamine in adult and adolescent mice and alterations in dopaminergic neurotransmission may be involved in this phenotype.

## Introduction

Tobacco smoking is a major public health issue and remains the single leading cause of preventable disease and death around the globe. Likewise, alcohol addiction is a pressing public health and socioeconomic concern. The use of each drug alone or in combination is the main preventable cause of premature death worldwide, with an estimated death toll of about 5 million individuals annually. Notably, tobacco use can lead to nicotine addiction and nicotine can serve as a gateway drug to facilitate intake of alcohol and other addictive drugs (Bechtholt and Mark 2002; DiFranza and Guerrera 1990; Horger et al. 1992; Hutchison and Riley 2008; Kandel and Kandel 2015; Kandel and Kandel 2014; Kelley and Rowan 2004; Kouri et al. 2001; Levine et al. 2011; Li et al. 2014; McQuown et al. 2007; McQuown et al. 2009; Meliska et al. 1995; Natividad et al. 2010; Rinker et al. 2011; Rosenberg 2014; Schindler et al. 2012; Schneider et al. 2012). In particular, nicotine has been reported to serve as a gateway drug for subsequent use and abuse of amphetamine, cocaine, and morphine (Baker et al. 2018; Baker et al. 2013; Bechtholt and Mark 2002; Gossop et al. 2006; Horger et al. 1992; Hutchison and Riley 2008; Kelley and Rowan 2004; Kouri et al. 2001; Levine et al. 2011; Li et al. 2014; Marks et al. 2015; McQuown et al. 2007; McQuown et al. 2009; Ruiz et al. 2018; Stinson et al. 2005; Storey et al. 2016).

Previous studies have shown that nicotine use is initiated during adolescent period and manifests the use and abuse of alcohol and other drugs (Best et al. 2000; Businelle et al. 2013; Carmody et al. 1985; DiFranza and Guerrera 1990; Lisha et al. 2014; McKee et al. 2011; Rimm et al. 1995; Romberger and Grant 2004; Torabi et al. 1993; York and Hirsch 1995). Interestingly, previous studies have shown that nicotine exposure during adolescence alters the aversive (Hutchison and Riley 2008) and rewarding (Kelley and Rowan 2004; Schochet et al. 2004) effects of cocaine. Additionally, it has been shown that aversion following nicotine withdrawal is reduced in adolescent than adults rats (O’Dell et al. 2007). Likewise, these authors have shown that female rats express enhanced nicotine-induced conditioned place preference (Torres et al. 2009), raising the possibility that adolescent and female adults are more prone to become polydrug users. Thus, in the present study we determined if the rewarding action of amphetamine would be altered by prior nicotine and alcohol exposure and this response would be different in adolescent (1-month-old) versus adult (2-3-month-old) male and female mice.

The neurotransmitter dopamine plays a critical role in the locomotor stimulatory and rewarding effects of amphetamine in dorsal and ventral striatum (Fleckenstein et al. 2007; Sulzer 2011). Previous studies have indicated the importance of dopamine transporter (DAT) in amphetamine mediated reward (Calipari et al. 2013). Amphetamine has been shown to regulate the release of dopamine via the reversal of DAT. Amphetamine enters cells through DAT or by passive membrane transport. Once inside the cell, amphetamine blocks monoamine oxidase, and alters vesicular storage of dopamine into synaptic vesicles. Consequently, accumulation of cytoplasmic dopamine leads to reversed transport of dopamine to extracellular space by DAT (Sulzer et al. 2005). Constitutive deletion of DAT expression in mice diminished amphetamine-induced accumbal dopamine release and associated hyperlocomotion, indicating the importance of DAT in these actions of amphetamine (Cagniard et al. 2014). Similarly, mice with overexpression of DAT compared to their wild-type controls displayed greater locomotor activity in response to amphetamine, had higher extracellular dopamine and showed preference to amphetamine at a much lower dose (Salahpour et al. 2008).

The NAc is an integral brain region in processing of reward. NAc receives dopaminergic input from ventral tegmental area (VTA) and glutamatergic input from prefrontal cortex (PFC), amygdala and hippocampus. Drugs of abuse, such as amphetamine, stimulate dopamine release in the NAc, leading to the activation of dopamine D_1_ or D_2_ receptors in the medium spiny neurons (MSNs). Therefore, we focused on the NAc and carried out Western Blot analyses to measure the level of DAT, dopamine D_1_ and D_2_ and glutamate receptors in this brain region. Our results showed that amphetamine induced a significant CPP response in control (exposed to saline previously) adolescent but not adult mice. However, mice of both age groups exhibited a comparable state-dependent CPP response following a challenge dose of amphetamine, showing that adolescent compared to adult control mice were more sensitive to the rewarding action of acute amphetamine under a drug-free but not drugged state. In contrast, the CPP response under both drug-free and drugged states was more robust in adult compared to adolescent mice with prior nicotine and alcohol experience. Correspondingly, DAT expression was greater in adult than young mice with prior nicotine and alcohol experience. Additionally, the level of D1 receptor was increased and that of D2 decreased in adult mice compared to adolescent mice with prior nicotine and alcohol experience. On the other hand, there were no significant changes in N-methyl-D-aspartate (NMDA) or α-amino-3-hydroxy-5-methyl-4-isoxazolepropionic acid (AMPA) receptors between mice of the two age groups.

## Materials and methods

### Animals

In this study we used young (one month of age), and adult (3-4 months of age) female and male mice (n = 7-8 mice of each age group) on a C57BL/6J mouse strain. Mice were bred in house and maintained 2-4 per cage with free access to laboratory chow and tap water and kept under a 12 h light/12 h dark cycle in a temperature- and humidity-controlled room. The light was on 6 AM and off at 6 PM. All experiments were conducted during the light cycle between the hours of 10:00 AM to 5:00 PM and were according to the National Institute of Health for the Proper Care and Use of Animals in Research and approved by the Institutional Animal Care and Use Committee at Western University of Health Sciences (Pomona, California, USA).

### Drug administration

Alcohol solutions were prepared from ethyl alcohol (200 Proof, OmniPur, EM Science; Gibbstown, NJ) in saline to a concentration of 0.2 g/ml, and injected via intraperitoneal (i.p.) route in a volume of 10 ml/kg body weight, yielding a dose of 2 g/ kg of ethanol. Nicotine free base from MP Biomedicals, Inc. (Solon, OH, USA) was dissolved in saline (0.025 mg/ml) and injected subcutaneously (s.c.) in a volume of 10 ml/kg body weight, yielding a dose of 0.25 mg/kg of nicotine free base. Amphetamine sulphate (Sigma/Aldrich) solutions (0.1 and 0.3 mg/ml) were prepared in saline and injected (i.p.) in a volume of 10 ml/kg body weight to yield doses of 1 and 3 mg/kg, respectively. The solutions were made daily prior to each conditioning and on the test day for state-dependent conditioned place preference (CPP).

### Experimental Procedures

#### Place Conditioning Paradigm

The place conditioning paradigm, widely used as a measure of preference and aversion (Bardo and Bevins 2000), was used to assess the motivational effects of nicotine and subsequent alcohol and amphetamine in adolescents and adult male and female mice. We used a 3-chambered place conditioning apparatus (ENV-3013, Med Associates Inc., St. Albans, VT). The details of the apparatus and place conditioning procedures are provided elsewhere (Nguyen et al. 2012; Tseng et al. 2013; Tseng et al. 2019). Briefly, on the preconditioning test day, mice were placed in the central neutral grey chamber of the apparatus and allowed to freely explore all the three chambers for 15 min under a drug-free state. The amount of time that mice spent in each chamber was recorded. Mice then received twice daily (morning and afternoon) conditioning (15 min each) for three consecutive days (on days 2-4) with nicotine/saline or saline/nicotine (0.25 mg/kg) or saline/saline and were then tested for place preference on day 5 (D5). The morning and afternoon conditionings were separated by 4 h. On days 6-8, mice received additional conditioning with nicotine (0.25 mg/kg) or saline, and then were tested for CPP 24 h later (D9). Mice conditioned with nicotine then received 3 conditioning with ethanol (2 g/kg) in the nicotine-paired chamber and saline in the saline-paired side and tested for CPP 24 h after the last alcohol conditioning (D13). On days 14-16, mice then received additional three conditioning with ethanol or saline as described above on days and were tested for CPP 24 h later (D17). Saline-conditioned control mice were injected with saline instead alcohol on the conditioning days and tested similarly. Finally, mice were conditioned (30 min each and once daily) with amphetamine (3 mg/kg) in the nicotine-alcohol-paired side and saline in the saline-paired chamber on days 18-19 and tested for CPP 24 h later (D20). The following day (D21), mice were tested for state dependent CPP, i.e., under a drugged state, in which mice were challenged with amphetamine (1 mg/kg) and then immediately tested for CPP. The injection order was counterbalanced such that mice received either drug or saline in the morning, and then the alternate treatment in the afternoon. On each postconditioning test day, mice were placed in the central chamber and allowed to freely explore the entire apparatus for 15 min. The amount of time that mice spent in each chamber was recorded in the same manner as for the preconditioning test day.

#### Western blot analysis

Immediately after completion of the last behavioural test, mice were deeply anesthetized with isoflurane (32%) and decapitated. Brains were quickly removed, frozen on dry ice and stored in a −80°C freezer. Nucleus accumbens (NAc) was dissected out and homogenized in RIPA lysis buffer (cat. # sc-24948, Santa Cruz) containing phosphatase and proteinase inhibitors. Protein concentration was estimated by Pierce™ BCA Protein Assay Kit (cat. # 23225). Protein lysates (45 ng) were prepared in equal volume with 2x Laemmli Sample Buffer (cat. #1610737, Bio-Rad) and loaded on 10% Mini-PROTEAN® TGX Precast Gel (Bio-Rad). The gel was transferred to PVDF membrane for 2 hours at 200mA. The membrane was then incubated with 1:1000 dilution of one of the primary antibodies, i.e., anti-mouse Actin, clone C4 (cat#MAB1501, Millipore), anti-rabbit dopamine D_2_ receptor (cat# AB5084P, Millipore), anti-rabbit AMPA receptor (GluR1) (D4N9V) (cat#13185, Cell Signalling), anti-rabbit NMDA receptor (GluN1) (D65B7) (cat# 5704, Cell Signalling); anti-rabbit dopamine transporter (cat# AB2231, Millipore); anti-rabbit dopamine D(1A) receptor (cat# ABN20, Millipore) overnight. The following day, the membranes were washed with tris-buffered saline (TBS) and 0.1% Tween 20three times and exposed to a secondary antibody (1:1000 dilution of anti-rabbit or anti-mouse) in a room temperature for 1 hour. The membrane then washed three times and Pierce™ ECL Western Blotting Substrate was added and bands were exposed using Bio-Rad ChemiDoc system. All antibodies were diluted in 1:1000 ratio, using 5% milk or 5% BSA for phosphorylated proteins, made in tris-buffered saline (TBS) and 0.1% Tween 20.

#### Data Analysis

Behavioural data are presented as mean (± S.E.M.) of the amount of time that mice spent in the drug-paired chamber (DPCh) and vehicle-paired chamber (VPCh) and analysed using three-way analysis of variance (ANOVA). Western blot data were analysed by two-way ANOVA. The Fisher’s post-hoc LSD test was used to reveal significant differences between adolescent vs. adult mice. A value of P < 0.05 was considered significant.

## RESULTS

### Amphetamine induced a significant CPP under a drug-free state in control adolescent but not adult mice

Figure 1 shows the amount of time that saline-treated control mice of each age group spent in the drug-paired (DPCh) and vehicle-paired (VPCh) chambers on days 1 (D1), 20 (D20) and 21 (D21). Three-way ANOVA revealed no significant effect of time (F_(2,156)_ = 1.689; P = 0.188), age (F_(2,156)_ = 0.233; P = 0.630), but there was a significant effect of context (F_(2,156)_ = 57.68; P<0.0001) and a significant interaction between time and context (F_(2,156)_ = 24.27; P<0.0001). The *post hoc* test revealed adolescent mice spent significantly more time in the amphetamine-paired chamber compared to saline-paired chamber, showing that these mice exhibited a robust CPP after conditioning (Fig. 1; right half of the graph; compared DPCh vs. VPCh on D20). On the other hand, conditioning with the same dose of amphetamine failed to induce CPP in control adult mice (Fig. 1; left half of the graph; compared DPCh vs. VPCh). When these mice were challenged with amphetamine (1 mg/kg) and tested under a drugged state the following day (D21), both adolescent and adult mice exhibited a significant state dependent and there was no difference in this response between adult and adolescent mice CPP (Fig. 1; compare DPCh vs. VPCH on D21 in each group).

**Fig. 1.**
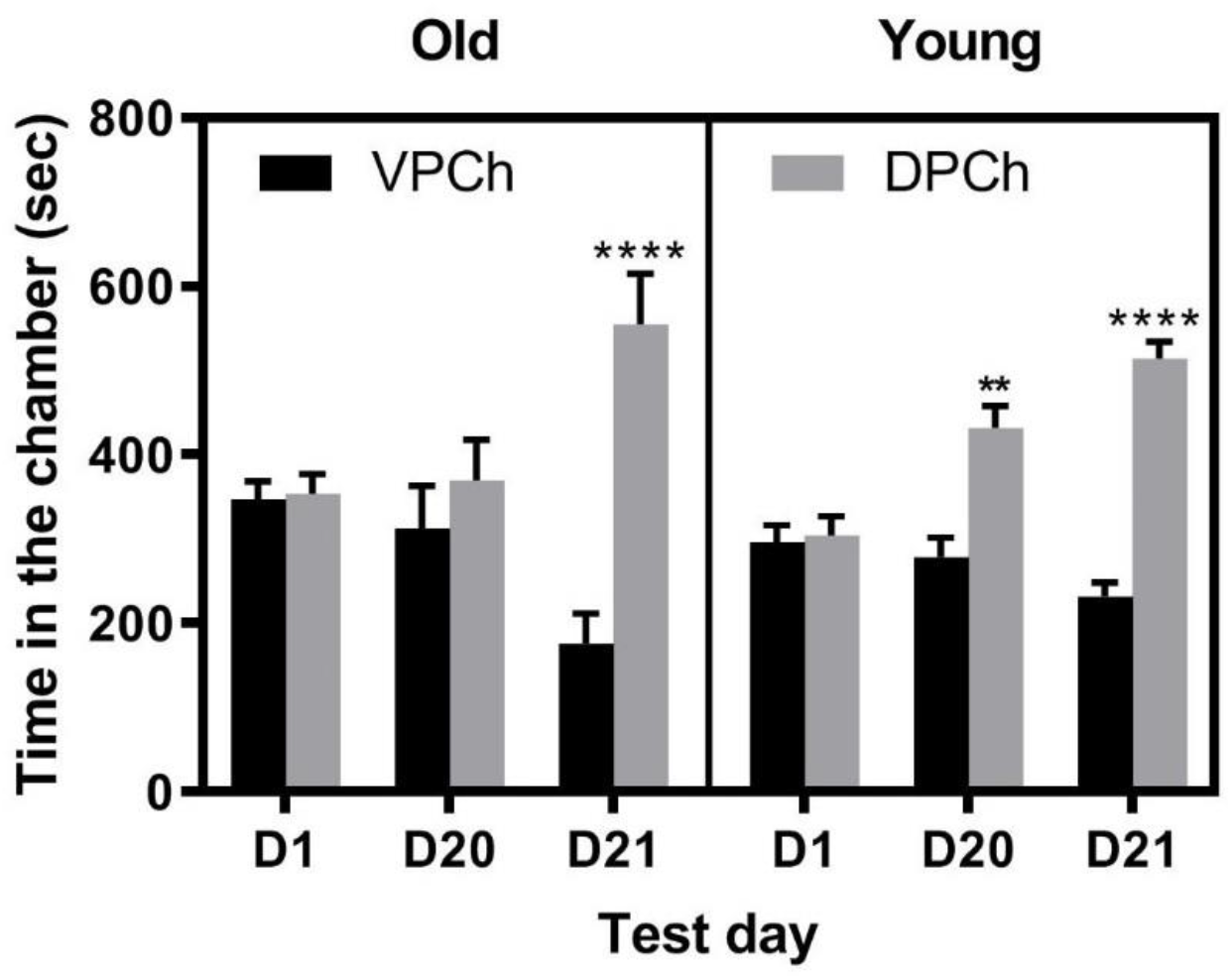
CPP induced by a single amphetamine conditioning in control adolescent and adult C57BL/6J mice. Male and female mice of both age groups (n = 7-8 mice per sex and age group) were tested for baseline and then received conditioning with saline in both place conditioning chambers. Mice were then conditioned with saline or amphetamine (3 mg/kg, i.p.) once daily in a counterbalanced manner and then tested for CPP 24 h after the last conditioning (D20). The following day, mice received a challenge dose of amphetamine (1 mg/kg) and tested for state dependent CPP (D21). Data represent the amount of time that mice spent in the drug-paired chamber (DPCh) or vehicle-paired chamber (VPCh) on the baseline test day (D1), CPP test day under a drug-free state (D20) and drugged state (D21) and analyzed by three-way ANOVA followed by the Fisher LSD test. **P<0.01, ****P<0.0001 vs. their respective VPCh

### Prior nicotine and alcohol conditioning facilitated the rewarding action of acute amphetamine under both drug-free and drugged states in adult compared to adolescent mice

Figure 2 illustrates the length of time that adolescent and adult mice with prior nicotine and alcohol exposure spent in the DPCh and VPCh prior to (D1) and after (D20) conditioning with amphetamine as well as on the test day for state dependent CPP (D21). Three-way ANOVA showed no significant effect of age (F_(2,168)_ = 1.135; P = 0.288), but there was a significant effect of time (F_(2,168)_ = 5.143; P = 0.007), a significant effect of context (F_(2,168)_ = 111.1; P<0.0001) and a significant interaction between time, age and context (F_(2,168)_ = 10.69; P<0.0001). The *post hoc* test revealed adult mice spent significantly more time in the amphetamine-paired chamber compared to saline-paired chamber on the CPP test under a drug-free state, showing that these mice exhibited a robust CPP response after conditioning with amphetamine (Fig. 2; left half of the graph; compared DPCh vs. VPCh on D20). On the other hand, conditioning with the same dose of amphetamine failed to induce CPP in adolescent mice with prior nicotine and alcohol experience (Fig. 2; right half of the graph; compared DPCh vs. VPCh). The challenge dose of amphetamine induced a significant state dependent CPP in mice of both groups, but the magnitude of this response was greater in adult than adolescent mice (P<0.0001).

**Fig. 2.**
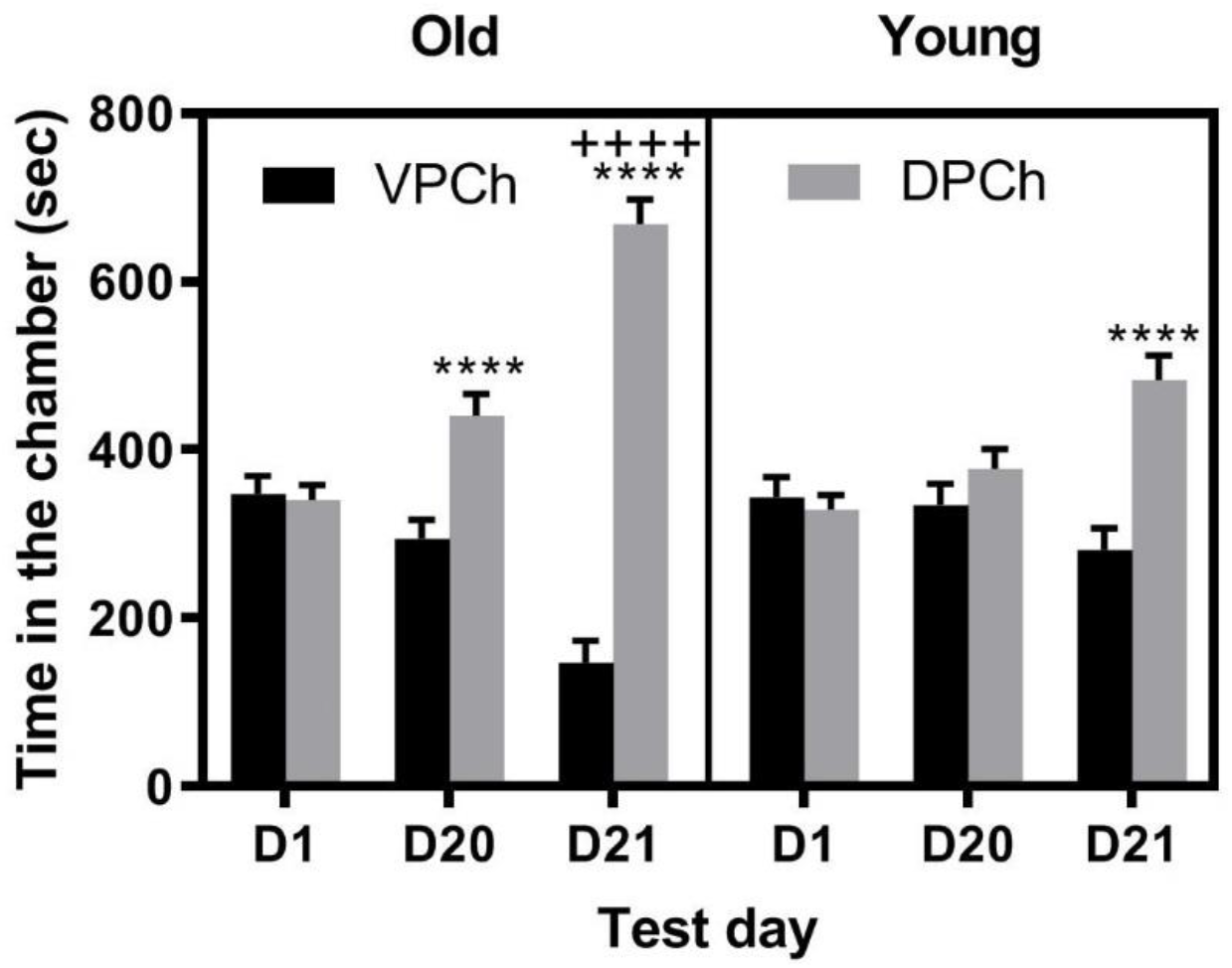
CPP induced by a single amphetamine conditioning in adolescent and adult C57BL/6J mice with prior nicotine and alcohol experience. Male and female mice of the two age groups (n = 7-8 mice per sex and age group) were tested for baseline and then received six conditioning with saline or nicotine (0.25 mg/kg, s.c.) followed by six conditioning with ethanol (2g/kg, i.p.) in the nicotine-paired chamber and saline in the saline-paired chamber. Mice were then conditioned with saline or amphetamine (3 mg/kg, i.p.) once daily in a counterbalanced manner and then tested for CPP 24 h after the last conditioning. Amphetamine conditioning was carried out in the nicotine-alcohol-paired chamber. Data represent the amount of time that mice spent in the drug-paired chamber (DPCh) or vehicle-paired chamber (VPCh) on the baseline test day (D1), CPP test day under a drug-free state (D20) and drugged state (D21) and analyzed by three-way ANOVA followed by the Fisher LSD test. ****P<0.0001 vs. their respective VPCh; ++++P<0.0001 vs. DPCh in Adolescents

### The expression of DAT and D_1_R was higher with concomitant decrease in D_2_R in adult compared to adolescent mice with prior nicotine and alcohol exposure

Given that the action of amphetamine is mediated via the reversal of dopamine transporter (DAT) and the increase in the release of dopamine (DA) in the synaptic terminal, we hypothesized that nicotine and alcohol may differentially alter the expression of DAT and/or dopamine D_1_ or D_2_ receptors (D_1_R D_2_R) in adult compared to adolescent mice. Consistent with this hypothesis, the expression of DAT in NAc was higher in adult compared to adolescent mice with prior nicotine treatment (Fig. 3). Furthermore, we observed a significant increase (P<0.05) in the expression of accumbal D_1_R with a concomitant decrease in the level of D_2_R in adult compared to adolescent mice with prior nicotine and alcohol treatment. To assess if these changes are selective to the dopaminergic neurotransmission, we also measured the expression of NMDA and AMPA (Fig. 3). Our results showed there was no change in the level of AMPA and NMDA receptors between adult and adolescent mice with prior nicotine treatment.

**Fig. 3.**
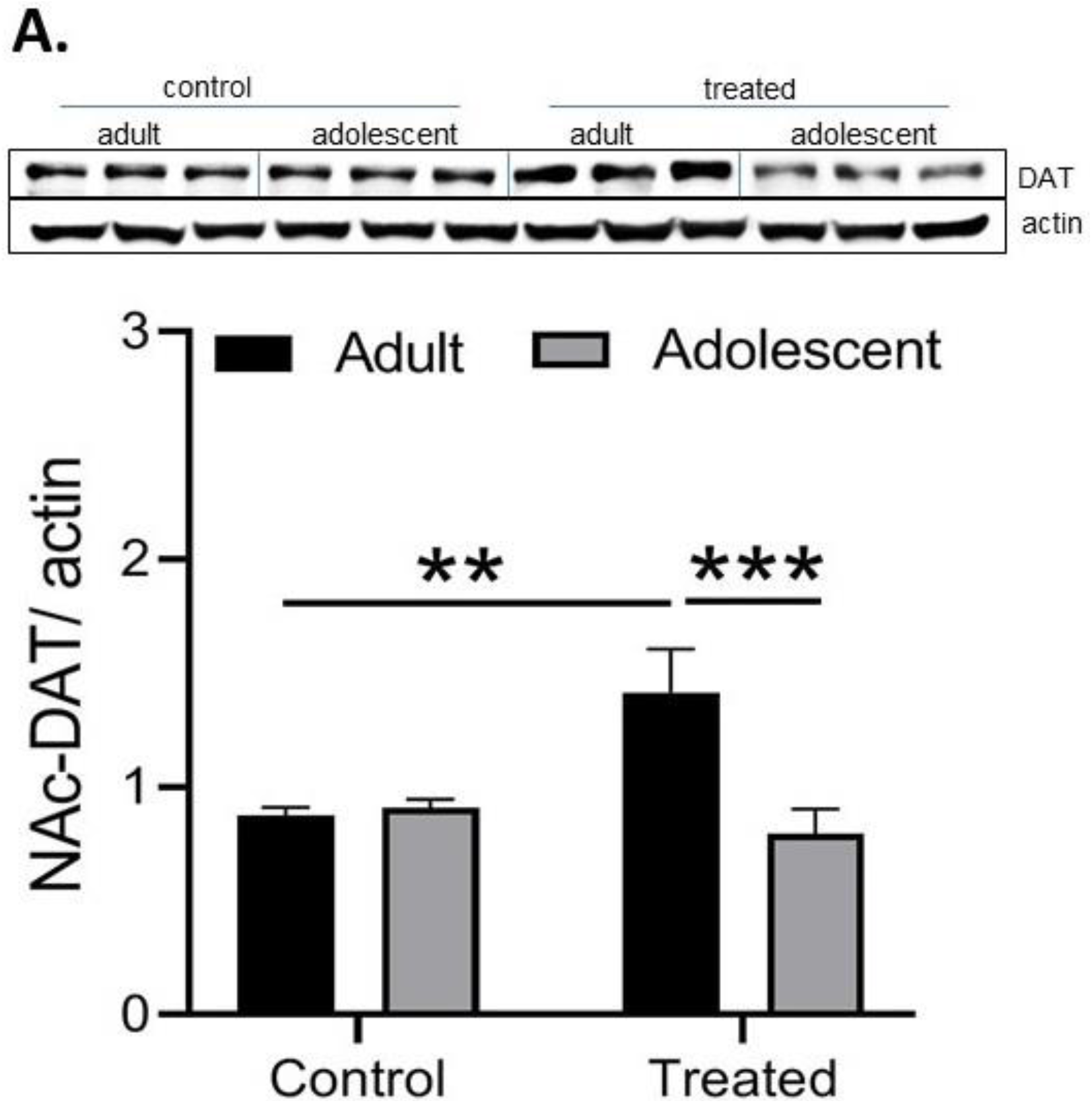
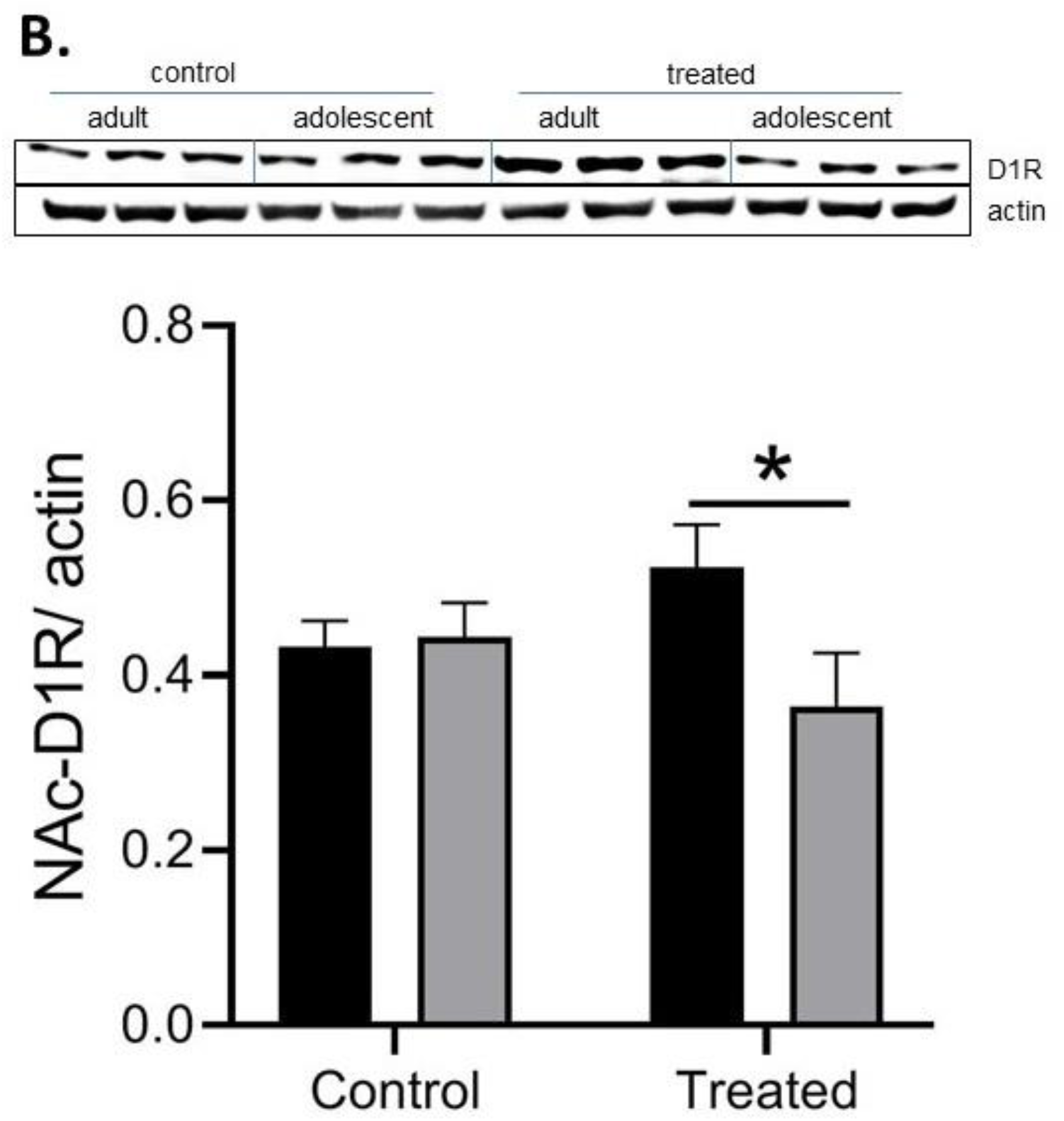
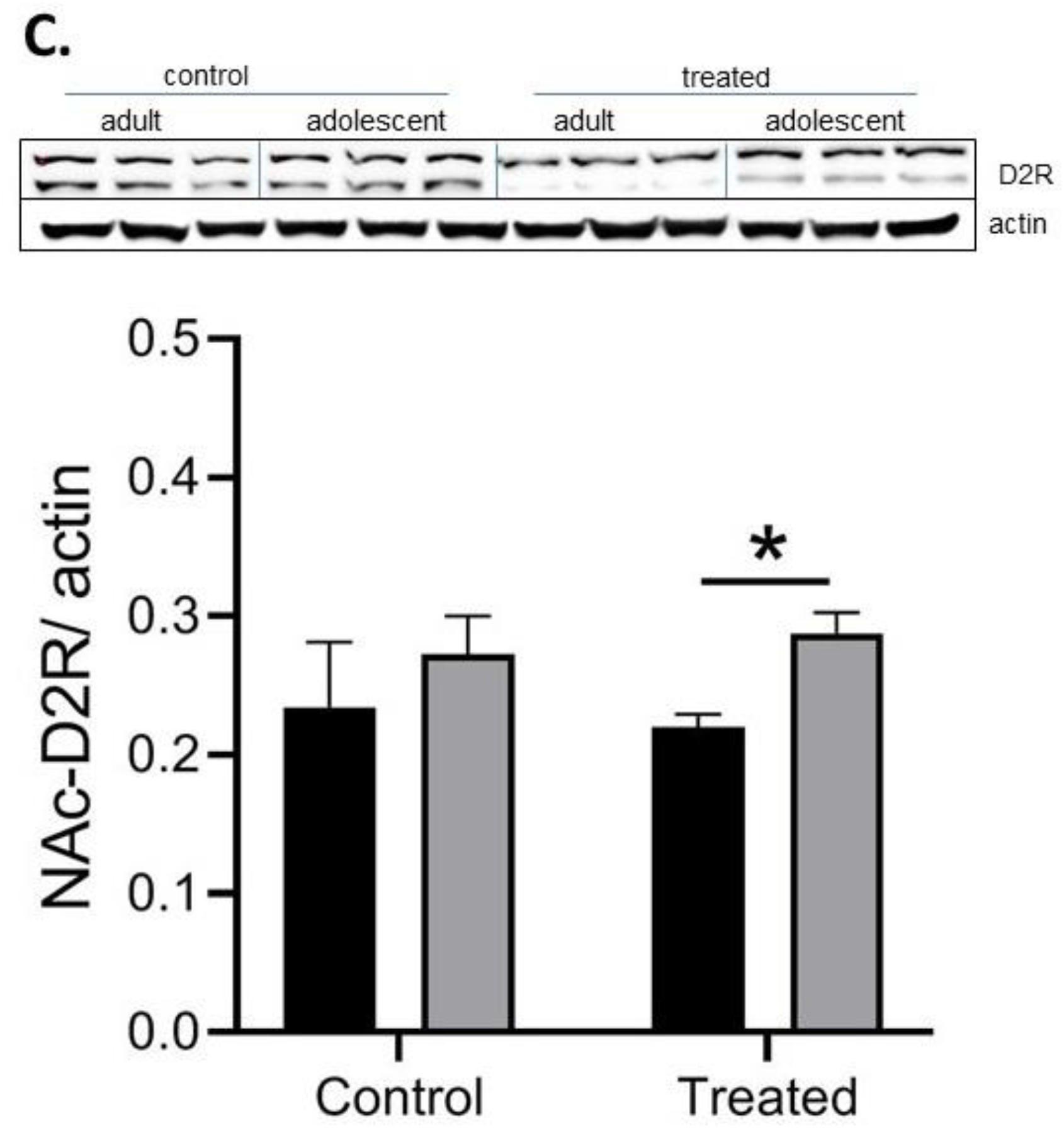
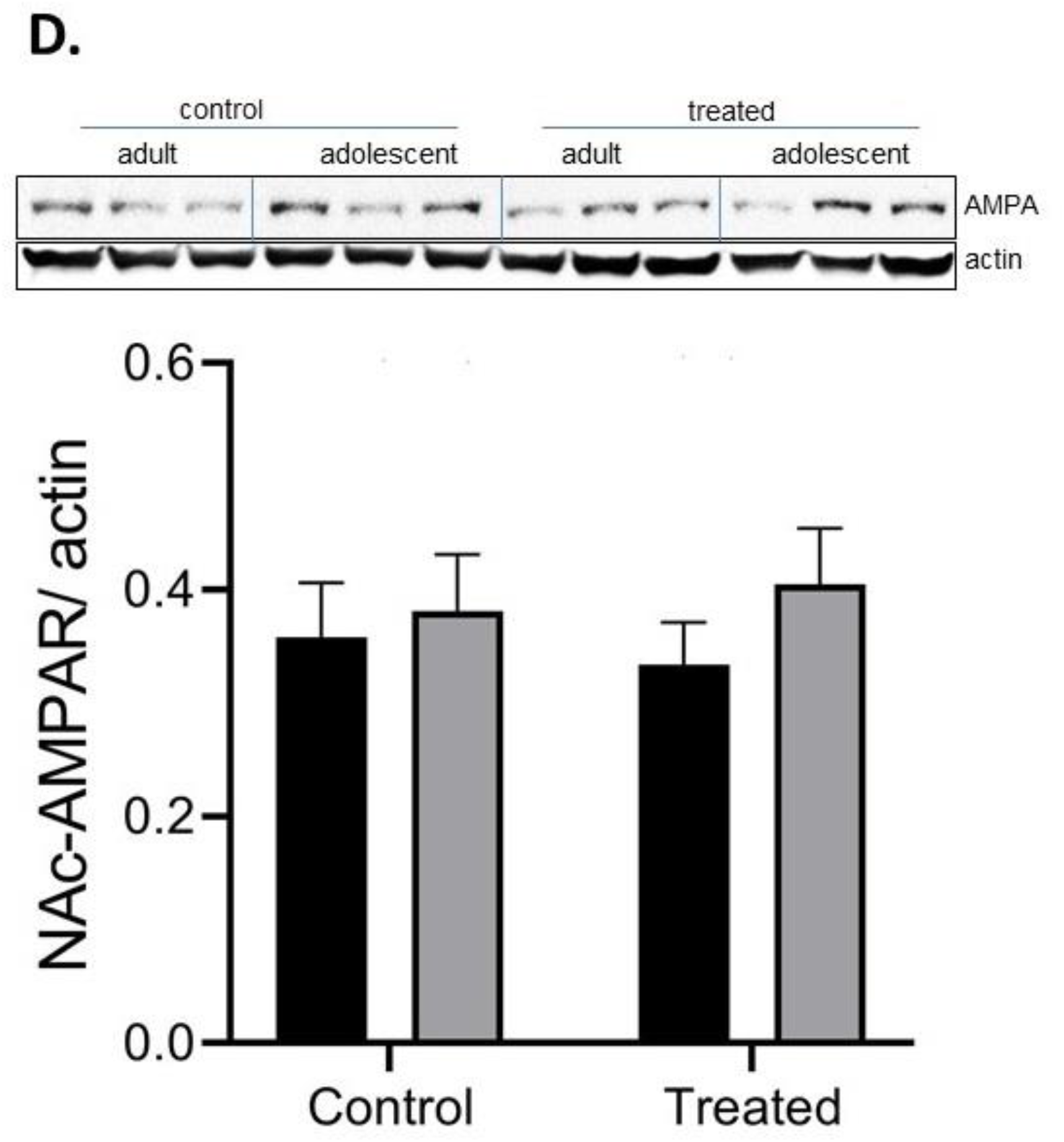
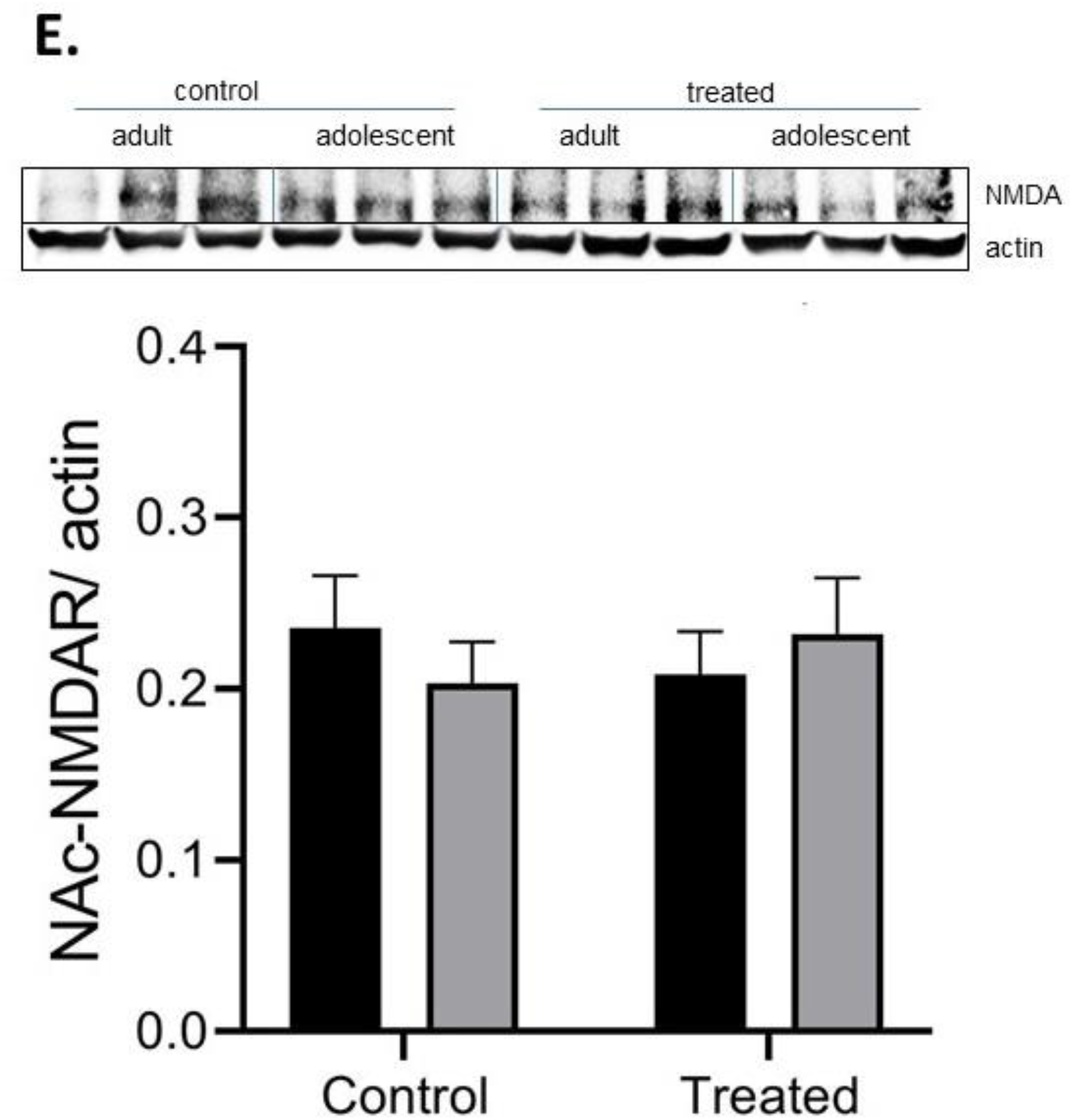
Expression of DAT and dopamine (D_1_ and D_2_) and glutamate (AMPA and NMDA) receptors in control and treated groups. Data represent the ratio of (**A**) dopamine transporter (DAT), (**B**) dopamine D1 receptors (D_1_R), (**C**) dopamine D2 receptors (D_2_R), (**D**) NMDA receptors and (**E**) AMPA receptors over beta-actin as well as their corresponding representative bands from adult male and female mice and were analyzed by two-way ANOVA, and then subjected to between-group comparisons using the Fisher’s least significant difference test. (**A**) **P<0.01 vs. control group, ***P<0.001 vs. treated adolescent group; (**B**) *P<0.05 vs. treated adolescent group; (**C**) *P<0.05 vs. treated adult group. Each band represents individual mouse (8-10 samples per treatment/ sex and each group)

## DISCUSSION

The main findings of the current study are that adolescent control mice exhibited a greater CPP response compared to their respective adult mice under a drug-free but not drugged state. On the other hand, prior nicotine and alcohol conditioning facilitated the rewarding action of amphetamine in adults but reduced it in adolescent mice under a drug-free state. Furthermore, the state dependent CPP was greater in adult compared to adolescent mice with prior nicotine and alcohol exposure. Along with these behavioural changes, we observed increases in the expression of DAT and D_1_R along with a reduction in D_2_R expression in adult compared to adolescent mice with prior nicotine and alcohol experience. Together, these results suggest that prior nicotine and alcohol treatment differentially affected the rewarding action of amphetamine in adolescent and adult mice, with concomitant changes in expression of proteins involved in dopaminergic neurotransmission.

Previous studies have shown differences in behavioural and neurochemical changes induced by amphetamines between adolescent and adult rodents (Baker et al. 2018; Ehrlich et al. 2002; Good et al. 2011). Consistent with these earlier studies, we found behavioural changes between control adolescent and adult mice following amphetamine conditioning. Our results showed that amphetamine induced a greater CPP response under a drug-free state in adolescent mice compared to adult mice. On the other hand, adolescent mice previously exposed to nicotine and alcohol showed reduced CPP following conditioning with the same dose of amphetamine. This is in contrast with previous studies showing that nicotine and alcohol each can serve as a gateway to heavier drugs. However, consistent with the gateway theory, we observed facilitation of amphetamine-induced CPP only in adult mice with prior nicotine and alcohol exposure compared to their saline-treated controls.

The CPP response under a drug-free state represents the motivational effect of the contextual cues that gain saliency following pairing with addictive drugs. It appears that the context gains more motivational valence following amphetamine pairing in adolescents than adult mice, which may explain the vulnerability of this population to use drugs after subsequent exposure to the same environment where they have used drug once, i.e., the context may serve as a stronger stimulus in facilitating drug use in this population. However, it appears that prior nicotine and alcohol exposure reduces this increased in motivational valence of the context in adolescent but increases it in adult mice, as we observed a robust CPP in adult mice with prior nicotine and alcohol experience but not in adolescent mice. The state-dependent CPP or the CPP response expressed under a drugged state, on the other hand, is a representative of the memory retrieval of the conditioned response that is acquired due to pairing of the subjective effects of the drug and the context during conditioning. This response is more robust than the response under the drug-free state in adult compared to adolescent mice with prior nicotine-alcohol exposure. Although we expected to observe a greater response in adolescent mice, it appears that prior nicotine and alcohol conditioning may differentially bring about molecular changes, thereby leading to these behavioural changes.

Substances of abuse, such nicotine, alcohol and amphetamine are thought to exert their rewarding effect through activation of mesocorticolimbic dopaminergic neurons (Pontieri et al. 1996). <u>Previous studies have shown that</u> deletion of DAT expression in mice diminishes amphetamine-induced accumbal dopamine release and associated hyperlocomotion (Cagniard et al. 2014). In contrast, mice with overexpression of DAT compared to their wild-type controls displayed greater locomotor activity in response to amphetamine, had higher extracellular dopamine and showed preference to amphetamine at a much lower dose (Salahpour et al. 2008). Given that we observed a greater amphetamine-induced CPP response under both drug-free and drugged states in adult mice with prior nicotine and alcohol exposure, we hypothesized that nicotine and alcohol may differentially alter the expression of DAT and/or dopamine D_1_ or D_2_ receptors (D_1_R D_2_R) in adult compared to adolescent mice. Consistent with this hypothesis we found that the expression of DAT and D_1_R was increased in adult mice with prior nicotine treatment compared to their respective adolescent mice as well as compared to control adult mice. Interestingly, there was a concomitant decrease in the expression of D_2_R in the NAc in adult mice with prior nicotine and alcohol experience compared to their respective adolescent mice. These changes were specific to the dopamine system because we did not observe changes in the expression of NMDA or AMPA receptors.

The observation that the expression of DAT and D_1_R increased and that of D_2_R decreased in adult mice with prior nicotine and alcohol exposure compared to adolescent mice may explain the underlying mechanism of the greater CPP response in adult mice compared to adolescent mice with prior nicotine and alcohol experience. We propose that the greater expression of DAT facilitates more amphetamine to get access to the presynaptic and once inside the cell and facilitates reversal of DAT, allowing more dopamine to exit through DAT and reaching the postsynaptic D_1_R. Considering that there is more D_1_R expression, there is more opportunity for dopamine to bind to postsynaptic D_1_R and elicits a greater CPP response (Fig. 4). Considering that there is a decrease in D_2_R expression in these mice compared to adolescent mice with prior nicotine and alcohol exposure, we expect a reduction in binding of dopamine to D_2_R and thus a decline in negative feedback mechanism, which also favours more dopaminergic neurotransmission to occur (Fig. 4). However, further studies are needed to delineate whether these changes are due to nicotine and/or alcohol exposure and/or combination of these drugs with amphetamine. Nevertheless, we do not believe that these alterations are due to amphetamine conditioning and/or amphetamine challenge alone since we did not observe these molecular changes between control adult and adolescent mice. Also, we do not believe that these changes are due to only nicotine exposure because adolescent rats have been shown to have lower basal levels of dopamine compared to adults in tissue samples of the striatum (Teicher et al. 1993) and reduced storage pool of releasable dopamine in this region (Stamford 1989). However, similar dopamine basal levels were found in tissue samples of NAc and frontal cortex between adolescent and adult rats (Teicher et al. 1993). Furthermore, nicotine has been shown to increase dopamine release in NAc in both adults and adolescent rats (Marusich et al. 2017). On the other hand, repeated ethanol administration in mice resulted in sensitization only in the adults, but not adolescents (Carrara-Nascimento et al. 2020). These authors also showed that adolescent mice exhibited lower dopamine levels in the PFC and NAc following ethanol exposure, compared to adults(Carrara-Nascimento et al. 2020). Cumulatively, based on the published data, it could be suggested that a greater CPP response following amphetamine in adult mice, observed in our study, could be related to dynamic changes in ethanol-induced decrease in dopamine release in NAc between adolescent and adult mice or a combination of action of nicotine and alcohol pre-exposure. Therefore, further studies are needed to assess the impact of preconditioning with each drug alone and their combination on the rewarding action of subsequent amphetamine treatment.

**Fig. 4.**
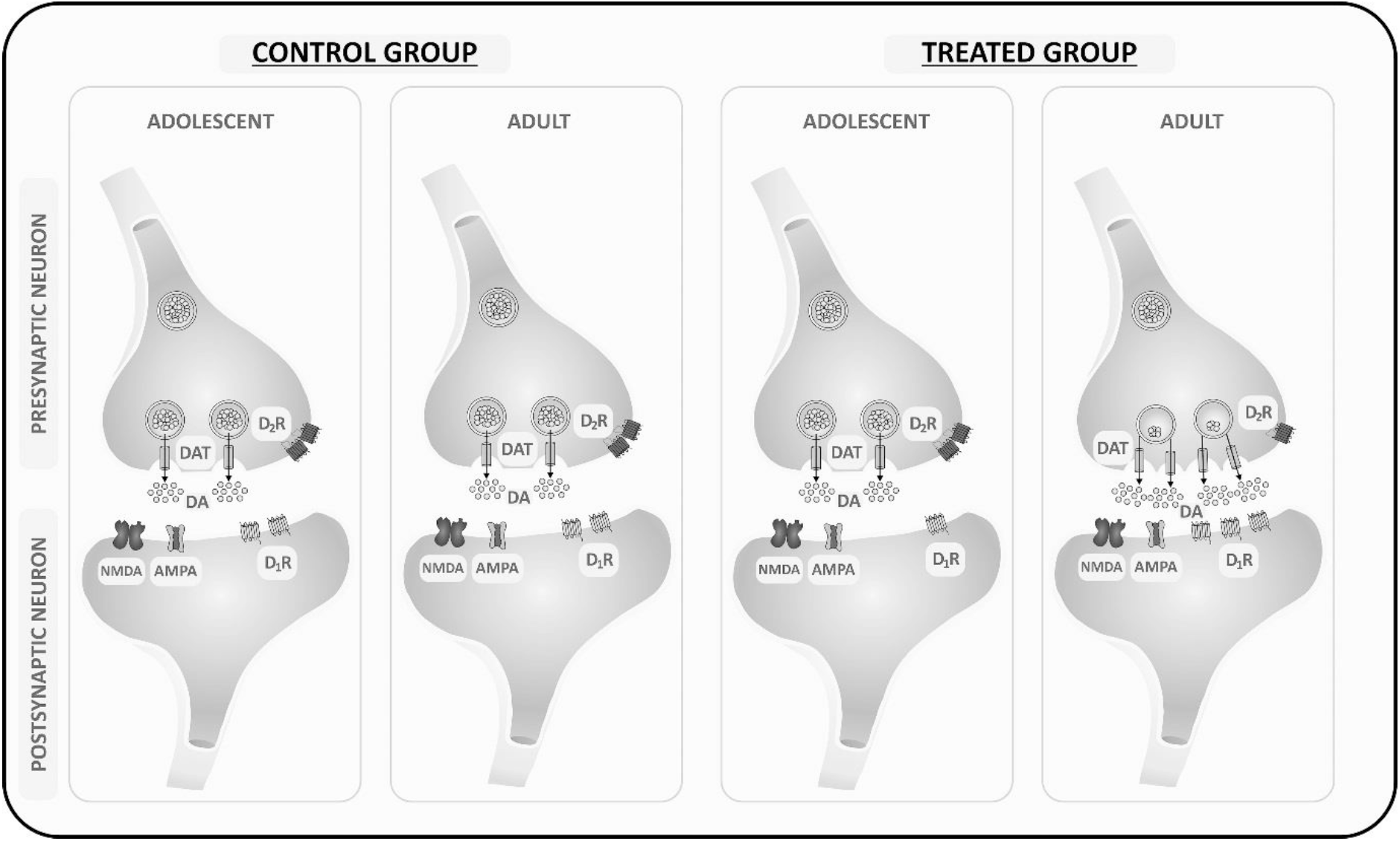
Schematic representation of changes in the level of DAT, D1 and D2 but not AMPA and NMDA receptors by prior nicotine and alcohol exposure, leading to altered amphetamine-induced stated-dependent CPP in adults vs. adolescent mice. Amphetamine induced a comparable CPP under a drugged state in control adolescent and adult mice. This may because the level of dopamine transporter (DAT) and dopamine (D_1_ and D_2_) receptors are not different between mice of the two age groups. On the other hand, the greater expression of DAT and D_1_ receptors and reduced levels of D_2_ receptors in adult than adolescent mice led to a greater CPP response under both drug-free and drugged states in adult than adolescent mice with prior nicotine and alcohol experience.

In summary, our results showed that amphetamine induced a greater reward in control (no prior drug experience) adolescent than adult mice but induced a comparable state dependent CPP in both age group. On the other hand, the rewarding action of amphetamine under both drug-free and drugged-state was reduced in adolescent and enhanced in adult mice with prior nicotine and alcohol exposure. These changes were associated with increased expression of DAT and D_1_ receptors and reduced level of D_2_ receptors. The decrease in the rewarding action of amphetamine in polydrug user adolescent compared to adult mice may suggest that this may be the cause for escalation of dose to chase pleasure in this vulnerable population.

## ACKNOWLEDGMENTS

The authors wish to thank Abdul Hamid and Osman Farhad for their assistance with the mouse breeding colony and preparing buffers, respectively. These studies were supported in part by Tobacco Related Disease Research Program (TRDRP) Grant # 24RT-0023H to KL.

## Compliance with ethical standards

### Conflict of interest

The authors reveal no conflict of interest.

### Data Sharing

Data will be made available upon the acceptance of this article from the corresponding author.

### Authors contribution

KL: study concept and design; AS: animal data acquisition; AS, SMA and KL: data analysis and interpretation; SMA and KL: Graphical Abstract and Diagram; AS: manuscript drafting; AS, SMA and KL: manuscript revision; all authors critically reviewed content and approved the final version for publication.

## Notes

### Competing Interest Statement

The authors have declared no competing interest.

